# Oncogenic KRAS alters splicing factor phosphorylation and alternative splicing in lung cancer

**DOI:** 10.1101/2022.05.20.492866

**Authors:** April Lo, Maria McSharry, Alice Berger

## Abstract

**Background:** Alternative RNA splicing is widely dysregulated in cancers including lung adenocarcinoma, where aberrant splicing events are frequently caused by somatic splice site mutations or somatic mutations of splicing factor genes. However, the majority of mis-splicing in cancers is unexplained by these known mechanisms. We hypothesize that the aberrant Ras signaling characteristic of lung cancers plays a role in promoting the alternative splicing observed in tumors.

**Methods:** We recently performed transcriptome and proteome profiling of human lung epithelial cells ectopically expressing oncogenic KRAS and another cancer-associated Ras GTPase, RIT1. Unbiased analysis of phosphoproteome data identified altered splicing factor phosphorylation in KRAS-mutant cells, so we performed differential alternative splicing analysis using rMATS to identify significantly altered isoforms in lung epithelial. To determine whether these isoforms were uniquely regulated by KRAS, we performed a large-scale splicing screen in which we generated over 300 unique RNA sequencing profiles of isogenic A549 lung adenocarcinoma cells ectopically expressing 75 different wild-type or variant alleles across 28 genes implicated in lung cancer.

**Results:** Mass spectrometry data showed widespread downregulation of splicing factor phosphorylation in lung epithelial cells expressing mutant KRAS compared to cells expressing wild-type KRAS. We observed alternative splicing in the same cells, with 2196 and 2416 skipped exon events in KRAS^G12V^ and KRAS^Q61H^ cells, respectively, 997 of which were shared (p < 0.001 by hypergeometric test). In the high-throughput splicing screen, mutant KRAS induced the greatest number of differential alternative splicing events, second only to the RNA binding protein RBM45 and its mutant allele RBM45^M126I^. We identified ten high confidence cassette exon events across multiple KRAS variants and cell lines. These included differential splicing of the Myc Associated Zinc Finger (MAZ). As MAZ regulates expression of KRAS, this splice variant may be a mechanism for the cell to modulate wild-type KRAS levels in the presence of oncogenic KRAS.

**Conclusion:** Proteomic and transcriptomic profiling of lung epithelial cells uncovered splicing factor phosphorylation and mRNA splicing events regulated by oncogenic KRAS. These data suggest that in addition to widespread transcriptional changes, Ras signaling pathways in cancer promote post-transcriptional splicing changes that may contribute to oncogenic processes.

## Background

Lung cancers are the leading cause of cancer death worldwide, with lung adenocarcinomas accounting for over 40% of lung cancers (American Cancer Society 2021). Lung adenocarcinomas, a form of non-small cell lung cancer, exhibit a high somatic mutational burden, with a median of 8.7 exonic somatic mutations per megabase in the tumor genome (Campbell et al. 2016). While most mutations are not known to have functional consequences, an important subset disrupts the pathways and molecular processes that regulate cell growth and proliferation, contributing to carcinogenesis.

One molecular process that is under a high level of regulation is RNA splicing and processing. RNA splicing is required in eukaryotic cells to produce functional proteins and to modulate protein abundance. In a phenomenon known as alternative splicing, cells can produce different versions of proteins from the same gene, contributing to the diversity and functional complexity of the human proteome. Analyses of cancer genomes and transcriptomes show that alternative splicing dysregulation is widespread in many cancers including in the lung (Kahles et al. 2018). One mechanism by which splicing can be altered in cancer is *cis*-acting splice site mutations which disrupt the recognition of exon splice sites (Jayasinghe et al. 2018). For instance, a mutation in a splice site of the tyrosine kinase gene *MET* leads to exon 14 skipping and a constitutively active protein (Lu et al. 2017; Onozato et al. 2009). Notably, lung cancers harboring this mutation confer clinical sensitivity to MET inhibitors (Frampton et al. 2015; Drilon et al. 2020). RNA splicing can also be altered in lung tumors by *trans*-acting mutations in splicing factor genes including *U2AF1, RBM10*, and *SF3B1* (Brooks et al. 2014; Zhao et al. 2017; Imielinski et al. 2012). Improved knowledge of these splicing aberrations and the resulting mis-spliced genes would improve discovery of new small molecule or gene therapies for cancer.

However, most aberrant splicing observed in cancer cannot be explained by known splice site or splicing factor mutations (Li et al. 2017). Another route to aberrant splicing is through dysregulation of signaling pathways upstream of splicing factors. Splicing factors, the proteins that determine where and when splicing occurs, are themselves regulated by intracellular signaling cascades (Zhou and Fu 2013). SR proteins are a family of splicing factors that plays a large role in splice site selection (Bradley et al. 2015). The function of SR proteins as recruiters of the spliceosome and catalyzers of the splicing reaction is heavily dependent on their phosphorylation state (Mermoud et al. 1992). As such, these proteins are regulated by kinases including SRPK1 and Clk/Sty (Shepard and Hertel 2009), and changes in their activity are associated with advanced forms of lung adenocarcinomas (Gout et al. 2012). In particular, previous studies show that alternative splicing can be disrupted through the AKT-SRPK-SR protein axis or the TGFb-CLK/sty-SR axis (Zhou et al. 2012; Nowak et al. 2008; Blaustein et al. 2005).

The AKT-SRPK signaling pathway is one of many arms of the receptor tyrosine kinase (RTK) and Ras signaling cascade, a network that regulates many cellular processes and is often altered in cancers (Mukhopadhyay et al. 2021). Mutations in *KRAS* are seen in ∼27% of lung adenocarcinomas, and in total, ∼76% of tumors harbor mutations in RTK-Ras-Raf pathway genes including the tyrosine kinases *EGFR* and *MET* (Imielinski et al. 2012; Cancer Genome Atlas Research Network 2014; Campbell et al. 2016). Despite a large body of work studying the Ras pathway, therapies targeting KRAS have only recently become viable and are currently effective only for the KRAS G12C variant (Canon et al. 2019). Inhibitors targeting EGFR or other RTKs are effective only for a limited time, with acquired resistance a nearly universal occurrence (Passaro et al. 2021; Drosten and Barbacid 2020). Understanding novel downstream processes of this oncogenic signaling cascade would expand the options for therapeutic intervention. In some instances, splicing factors or alternative isoforms can be amenable to inhibition by small molecule inhibitors, providing opportunities to relieve tumor burden (da Silva et al. 2015). Moreover, oligonucleotide therapies for aberrant splicing have proven effective in treating genetic disorders and could be used in precision oncology as well (Finkel et al. 2016).

In this study, we ask which mRNA splicing processes are downstream of and directed by Ras signaling. We employ a controlled experimental system of lung cell lines in order to investigate the effects of specific genetic perturbations on alternative splicing. In doing so, we avoid confounding influences of other contextual differences on splicing and increase the sensitivity to uncover oncogene-regulated aberrant splicing events.

## Results

### Mutant KRAS suppresses splicing factor phosphorylation in lung epithelial cells

The process of mRNA splicing is known to involve regulation of phosphorylation states (Cho et al. 2011; Aubol et al. 2016). Previously, we generated human AALE lung epithelial cells stably expressing *KRAS*^*WT*^, *KRAS*^*G12V*^, *KRAS*^*Q61H*^, *RIT1*^*WT*^, and *RIT1*^*M90I*^, and performed liquid chromatography and tandem mass spectrometry (LC-MS/MS) to profile their proteomes and phosphoproteomes (Lo et al. 2021). As expected, phosphorylation of Ras pathway proteins was significantly altered in cells expressing KRAS or RIT1 variants (Fig. 1A). Interestingly, in cells expressing KRAS^G12V^ or KRAS^Q61H^ variants, we also observed a marked change in the phosphorylation of splicing factor proteins (Fig. 1A) with several SR proteins showing decreased phosphorylation state (Fig. 1B). As each phosphosite was normalized to total protein level, and total protein level of these SR proteins was unchanged (Supplemental Fig. 1A), the decreased SR protein phosphorylation was likely due to differences in phosphorylation state itself. The phosphosites that showed the greatest decrease in phosphorylation levels (Fig. 1B) occurred in proteins known to be precisely regulated by phosphorylation: SRSF7, SRSF1, and SRSF2 (Cho et al. 2011; Aubol et al. 2003; Naro and Sette 2013). Notably, alteration of SRSF7, SRSF1, and SRSF2 protein phosphorylation occurred predominantly in KRAS^mut^ cells compared to KRAS^WT^ cells, and not in the RIT1^M90I^ cells compared to RIT1^WT^ cells (Fig. 1C, Supplemental Fig. 1B). These data suggest that KRAS may uniquely alter SR protein phosphorylation state in a manner distinct from another RAS-family GTPase, RIT1.

**Figure 1.**
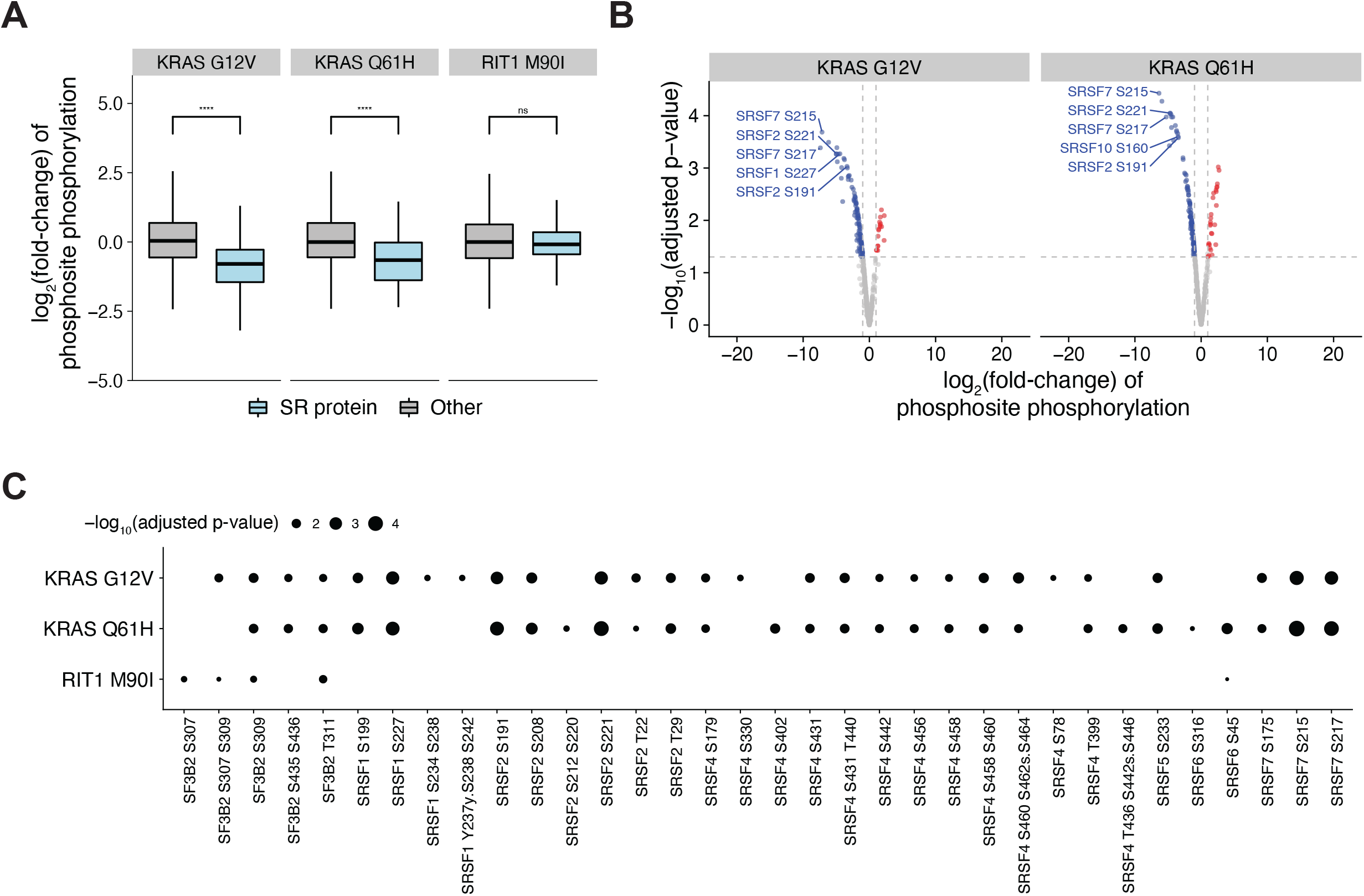
Mutant KRAS suppresses splicing factor phosphorylation in lung epithelial cells. A) Phosphorylation changes in SR proteins (blue) or other proteins (gray) in KRAS^G12V^ or KRAS^Q61H^ vs KRAS^WT^ cells, and RIT1^M90I^ vs RIT1^WT^ cells; * p<0.05, **** p<0.0001 by one-sided Wilcoxon rank-sum test. B) Volcano plots of the phosphorylation changes in mRNA splicing proteins. Sites with less phosphorylation in KRAS^G12V^ or KRAS^Q61H^ compared to KRAS^WT^ are shown in blue, sites with more phosphorylation are shown in red. The most significantly altered SR protein phosphosites are labeled. C) Heatmap of phosphorylation changes in phosphosites of RNA splicing proteins of interest SF3B2, SRSF1, SRSF2, SRSF4, and SRSF7.

### Oncogenic KRAS regulates alternative splicing in lung epithelial cells

Following the observation that splicing factors are differentially phosphorylated in KRAS^mut^ overexpressing cells compared to KRAS^WT^ overexpressing cells, we sought to directly compare RNA splicing regulated by oncogenic KRAS. To this end, we performed RNA sequencing and differential splicing analysis using rMATS (Shen et al. 2014). Alternative splicing of cassette exons (Fig. 2A) is tightly regulated by SR proteins (Zhou and Fu 2013; Zheng et al. 2020) and these events are typically the most prevalent form of alternative splicing detected in cancer and development (Scotti and Swanson 2016; Singh and Cooper 2012). The majority of splicing events observed in AALE cells were cassette exon events (70.3%), so we focused on cassette exon events for subsequent analyses. We observed that many cassette exons are more skipped or more promoted in KRAS^G12V^ and KRAS^Q61H^ cells compared to KRAS^WT^ cells (Fig. 2B). In addition, 997 exons are differentially skipped in both KRAS^G12V^ and KRAS^Q61H^ cells compared to the wild-type cells, indicating that oncogenic KRAS variants cause similar changes in splicing patterns (hypergeometric test for intersect p < 0.0001) (Fig. 2C). The exons most differentially expressed in both KRAS^mut^ cell lines include exons *MOK*, which encodes a member of the MAP kinase superfamily, and *NT5C2*, which encodes a 5′-nucleotidase enzyme implicated in chemotherapy resistance (Qian et al. 2015; Tzoneva et al. 2013) (Fig. 2D). These shared cassette exon events point to a change in splicing regulation driven by activated KRAS.

**Figure 2.**
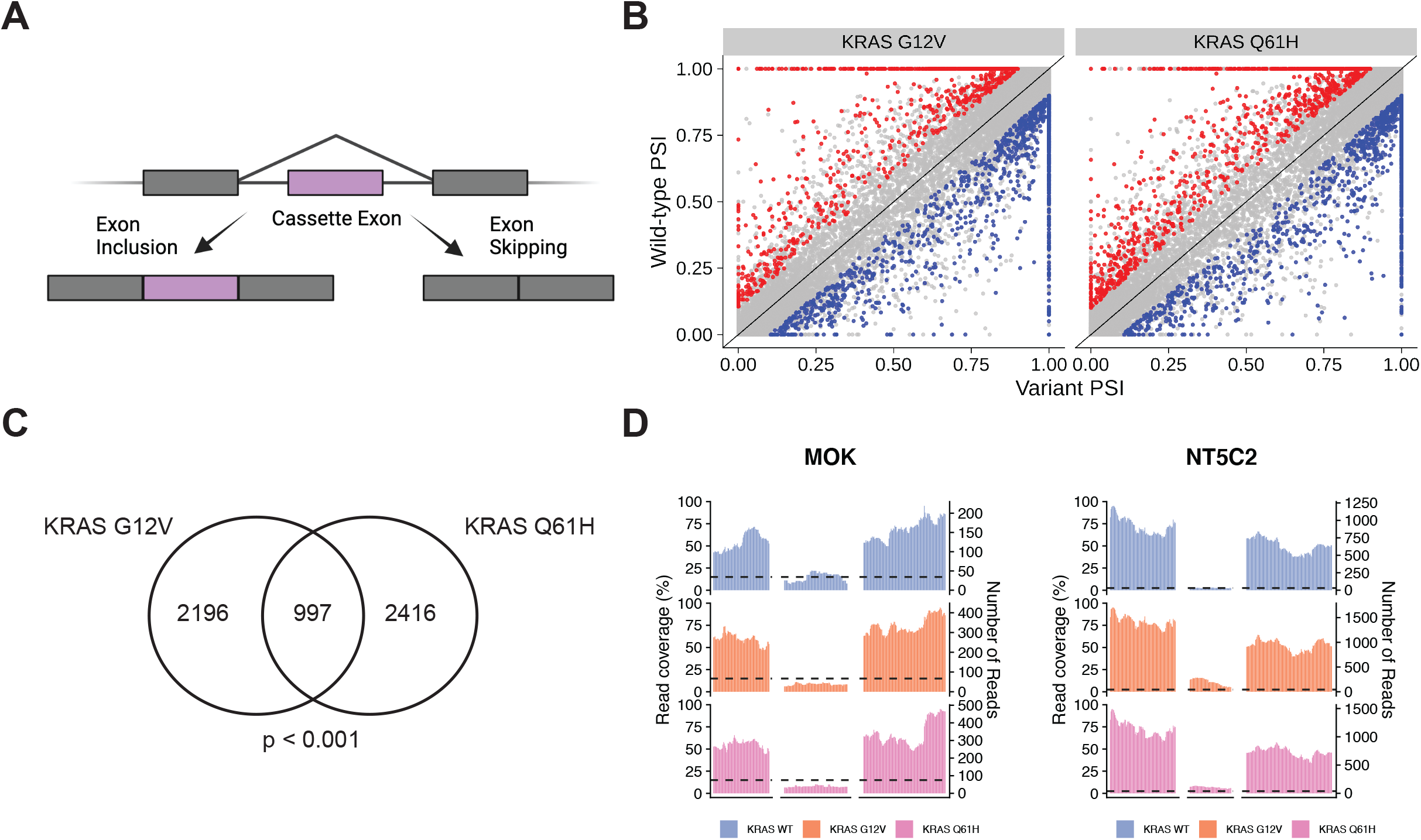
Oncogenic KRAS regulates alternative splicing in lung epithelial cell. A) Schematic of the inclusion or skipping events of a cassette exon. B) Differentially spliced exons in KRAS^G12V^ and KRAS^Q61H^ cells compared to KRAS^WT^ cells. PSI = Percent Spliced In. Red = exons promoted at least 10% more in KRAS^WT^ than KRAS^mut^. Blue = exons promoted at least 10% more in KRAS^mut^ than KRAS^WT^. FDR < 0.05. C) Overlap of differentially spliced exons in KRAS^G12V^ vs KRAS^WT^ and KRAS^Q61H^ vs KRAS^WT^. P-value calculated by modeling skipped exon events as a hypergeometric distribution. D) Read coverage plots of skipped exon events in *MOK* and *NT5C2*, normalized by total RNA-seq library size and RNA composition. Left axis = percent coverage relative to region. Right axis = absolute read coverage.

### A large-scale transcriptome screen as a platform for splicing discovery

To further study how KRAS and related oncogenes regulate alternative splicing, we performed a large scale perturbation screen in A549 lung adenocarcinoma cells. Isogenic A549 cell lines were generated by lentiviral transduction in 384 well format as described previously (Berger et al. 2016) with multiple biological replicates per variant. 4 days after transduction, cells were lysed for transcriptome analysis using the SmartSeq low input method (Fig. 3A). To identify oncogene-driven effects on transcription and splicing, we chose 28 genes to study based on high mutation frequency in lung adenocarcinomas and relevance in RNA processing and signaling pathways (Berger et al. 2016). In total, we generated whole transcriptome profiles for 374 unique replicates of arrayed isogenic cell lines expressing 79 different lentiviral constructs which include vectors for wild-type genes, variant alleles, and controls. For each replicate used in subsequent analyses, we confirmed overexpression of the allele by direct analysis of the RNA-seq data (Supplemental Fig. 3A-E).

**Figure 3.**
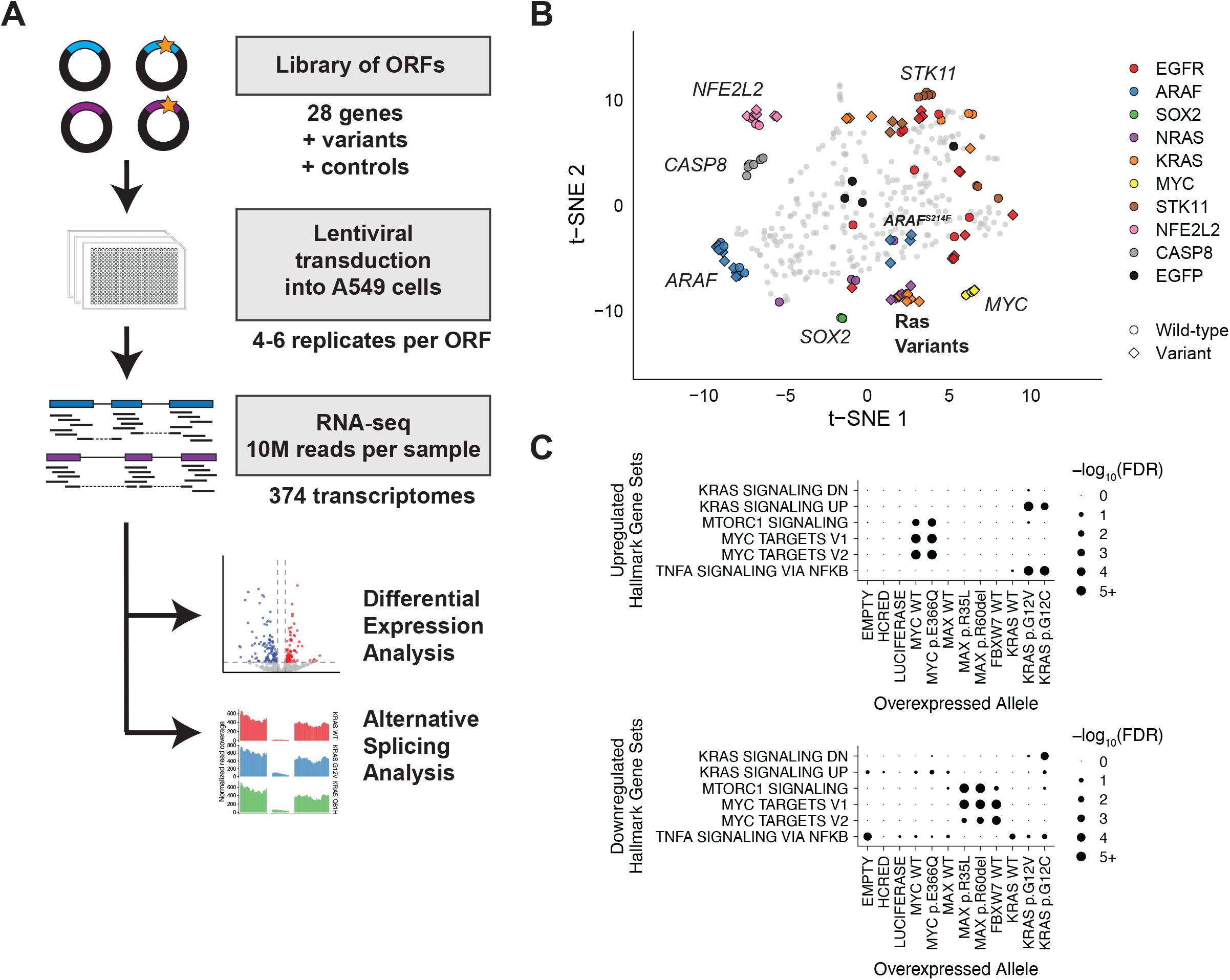
A large-scale transcriptome screen as a platform for splicing discovery. A) Experimental workflow for RNA-seq analysis of lung adenocarcinoma genes and variants. B) Transcriptome profiles of cells expressing different gene constructs. Dimensionality reduction performed using t-distributed stochastic neighbor embedding (t-SNE) (Van Der Maaten and Hinton 2008). Each point represents an experimental replicate. Genes and alleles of particular interest are colored and labeled. Controls are circles colored in black. Circle = wild-type allele. Diamond = variant allele. C) Gene set analysis with GOseq of up-or down-regulated transcripts in MYC, KRAS, FBXW7, and negative control cells. Hallmark gene sets from mSigDB.

Comparing gene expression signatures across the full experiment highlighted the distinct transcriptomic effects due to the ectopic expression of different open reading frames (ORFs). For example, ectopic expression of transcription factors such as *MYC, SOX2*, and *NFE2L2* resulted in consistent expression shifts across the transcriptome which are reflected by replicates which group close together (Fig. 3B). Expression profiles also highlighted the functional differences between wild-type and variant alleles of oncogenes. Consistent with previous findings, cells with activating ARAF^S214F^ showed similarities to *KRAS* and *NRAS* variant cells, while wild-type *ARAF*, neutral variant ARAF^V145L^, and the kinase-inactivated variants ARAF^S214F/D429A^ and ARAF^S214C*/*D429A^ did not (Berger et al. 2016).

To determine how ectopic expression of each gene affected the transcriptome, we inspected the differentially expressed genes between cell lines expressing an experimental ORF and cell lines expressing a control vector. As expected, overexpression of the transcription factor *MYC* induced high levels of *MYC* expression (Supplemental Fig. 3F). Gene set analysis further showed that the transcripts with increased expression in *MYC*-overexpressing cells include known MYC target genes (Fig. 3C). Conversely, overexpression of known MYC inhibitors such as *FBXW7* decreased the expression of MYC target genes. Expression of two inactivating *MAX* mutations, but not wild-type *MAX*, also suppressed *MYC* target gene expression (Yeh et al. 2018; Augert et al. 2020) (Fig. 3C). These findings demonstrate that this large-scale gene expression profiling screen can recapitulate known biology and support the utility of these data for further discovery.

One potentially confounding factor for this system is the natural endogenous KRAS^G12S^ variant found in A549 cells. To verify in this background that we were observing the activity of activated KRAS from ectopic expression of KRAS variants, we first ensured that KRAS was overexpressed by a significant amount. We defined significant allele overexpression as two standard deviations above endogenous KRAS expression (Supplemental Fig. 3A). Then we performed gene set analysis on A549 cells ectopically expressing KRAS^WT^, KRAS^G12V^, or KRAS^G12C^. Genes involved in upregulation of KRAS signaling were enriched in KRAS^G12V^ and KRAS^G12C^ cells but not KRAS^WT^ cells. Similarly, another gene set we saw enriched in KRAS^mut^ cells was the Nf-kb signaling pathway (Fig. 3C), which oncogenic KRAS is known to activate (Barbie et al. 2009; Daniluk et al. 2012). These and previous data suggest that despite their endogenous *KRAS* mutation, A549 cells can be used as a model for KRAS signaling upon further KRAS perturbation (Berger et al. 2016).

### A screen for splicing alterations in lung cancer identifies alternative splicing events regulated by KRAS

The use of RNA sequencing rather than probe-based array or bead technologies for transcriptome analysis provides the opportunity to measure alternative splicing in addition to differential gene expression. In both the data from AALE cells and the large-scale screen in A549 cells, we performed differential splicing analysis to first compare replicates with ORF expression against replicates expressing a control vector. We observed a mean of 456 differentially spliced events per overexpressed wild-type or variant allele and fewer than 1000 events in all except wild-type RBM45, which induced 1496 total differential alternative splicing events compared to vector control (Supplemental Fig. 5A-B, z-score = 4.51). Overexpression of the cancer-associated RBM45^D434Y^ allele also induced more alternative splicing events than average (z-score = 1.53).

Notably, several of the ORFs that perturbed splicing the most were transcription factors. These included *NFE2L2, SOX2*, or *MYC* (Supplemental Fig. 5A). Cells overexpressing wild-type *NFE2L2* harbored 931 alternative splicing events compared to vector controls (z-score = 2.06). Similarly, cells with wild-type *MYC* or the activated *MYC*^*E366Q*^ variant exhibited 787 and 726 differential splicing events, respectively (z-scores = 1.44 and 1.17). These increases may reflect a regulatory role played by mRNA splicing to adjust protein abundance levels when transcription activity and pre-mRNA levels are disrupted (Han et al. 2017).

We next compared cells overexpressing each variant allele to cells overexpressing the corresponding wild-type allele. Again, we observed the highest number of differential splicing events in RBM45 variant cells compared to RBM45 wild-type cells (z-score = 2.94, Fig. 4A). Besides RBM45, the variant with the greatest alternative splicing effect compared to the wild-type gene was KRAS^G12C^ with 729 total differential splicing events including 440 cassette exons. Similarly, KRAS^G12V^ cells differentially splice 409 events compared to KRAS^WT^ cells including 232 cassette exons (Fig. 4A). Since G12V, G12C, and Q61H are all KRAS activating variants that promote deregulated downstream MAPK signaling (Cook et al. 2021), we took advantage of our parallel analyses of multiple KRAS variants across two cellular contexts to identify cassette exon splice variants that are similarly differentially regulated in all four KRAS^mut^ AALE and A549 cell lines compared to the respective KRAS^WT^ cells. Using this approach, we identified 10 high-confidence alternative cassette exons that are consistently altered when KRAS is activated across the two different cellular contexts and three KRAS variants (Fig. 4B). The approximate degree of splicing change (percent spliced in) induced by KRAS^G12V^ was similar in A549 and AALE cells (Fig. 4B; Pearson’s coefficient = 0.71, p-value = 0.020).

**Figure 4.**
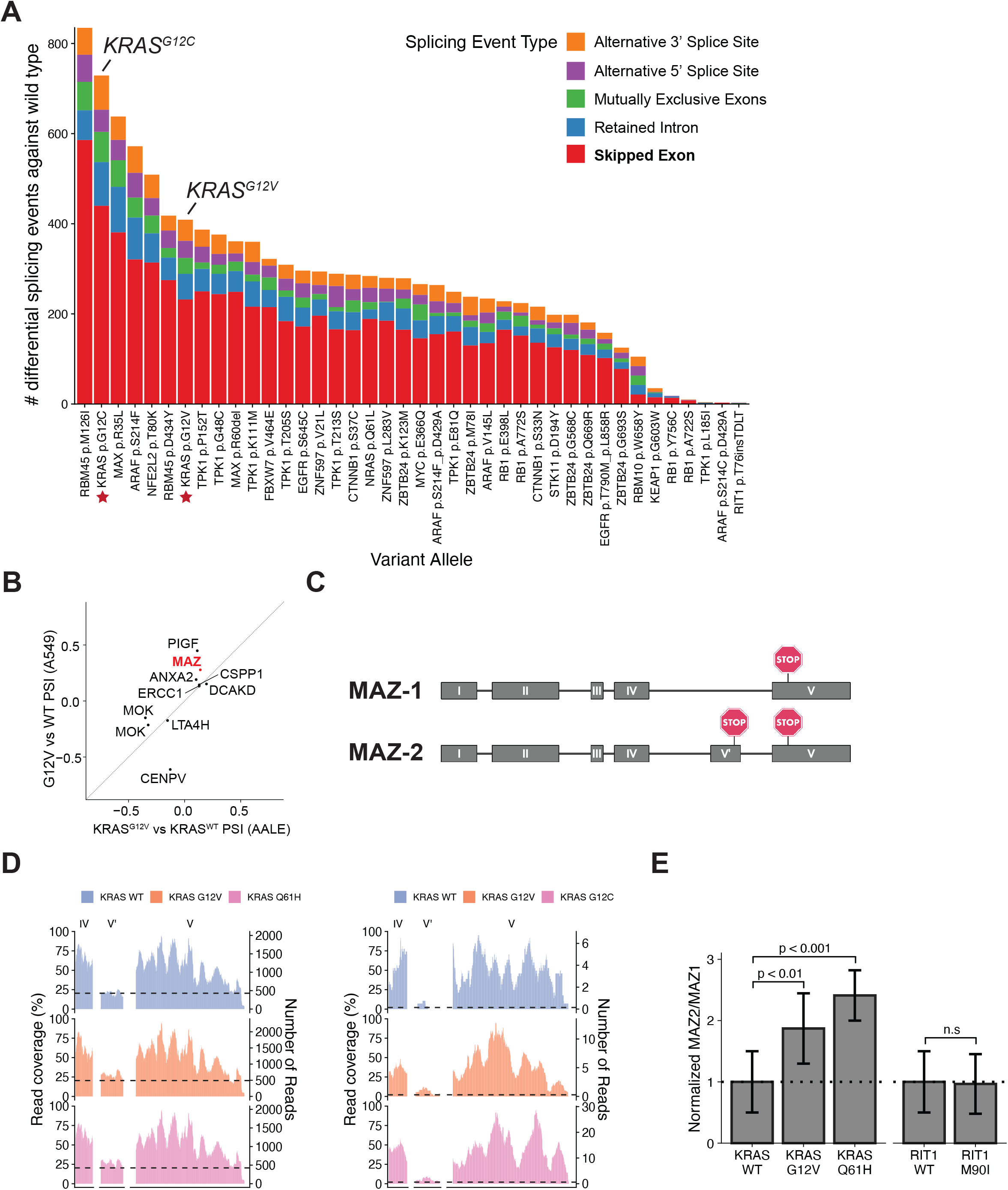
A screen for splicing alterations in lung cancer identifies alternative splicing events regulated by KRAS. A) Number of events differentially spliced between cells with variant alleles and cells with respective wild-type alleles. KRAS^G12C^ and KRAS^G12V^ are starred and labeled. Colors represent the five splicing event categories considered. FDR < 0.05. ΔPSI > 10%. B) Change in exon levels in cassette exons differentially expressed in both KRAS^G12V^ compared to KRAS^WT^ and KRAS^G12C^ compared to KRAS^WT^ R7M=^WT^, KRAS^G12V^, and KRAS^G12C^ overexpressing A549 cells (left) and KRAS^WT^, KRAS^G12V^, and KRAS^Q61H^ overexpressing AALE cells (right), relative to the expression of flanking exons. Inset zooms in on cassette exon. E) Relative levels of MAZ-2 compared to MAZ-1 isoforms in AALE cells overexpressing KRAS or RIT1 alleles, normalized to KRAS^WT^ or RIT1^WT^, as measured by quantitative RT-PCR. Error bars = standard error.

As our phosphoproteomic analysis indicated changes in the activity of SR proteins, we asked if these genes are differentially expressed. In most cells, including KRAS and EGFR overexpressing cells, there was no notable change in the gene expression of splicing factors such as *SF3B1, SRSF1, SRSF2*, or *SRSF7* (Supplemental Fig. 4C). We thus conclude that, in our experimental system and with our driver genes of interest, splicing factors are not substantially regulated at the mRNA level, and their contribution to the alternative splicing changes we observe are phosphorylation based.

Of the ten splice variants identified, we focused on the isoforms of the transcription factor Myc-associated zinc finger protein (MAZ) due to its previously characterized isoforms and its role in increasing the expression of *KRAS* and *HRAS* (Yang et al. 2019). The major mRNA isoform MAZ-1 is composed of five exons whereas isoform MAZ-2 has six exons, with the inclusion of exon V’ (Fig. 4C). This additional exon includes an early stop codon, leading to a protein isoform with an alternative C terminus (Ray et al. 2002). In both AALE and A549 cells, expression of KRAS^mut^ resulted in an increase in exon V’ inclusion (ΔPSI = .14 to .34, FDR < .01) (Fig. 4D). These alternatively spliced isoforms were validated through quantitative RT-PCR showing a lower MAZ-2 to MAZ-1 isoform ratio in KRAS^WT^ overexpressing AALE cells (Fig. 4E, Supplemental Fig. 5D). Thus, we found that KRAS^WT^ and KRAS^mut^ overexpressing cells display distinct MAZ isoform levels, suggesting that alternative splicing of the *MAZ* gene is uniquely regulated downstream of KRAS (Supplemental Fig. 5E).

## Discussion

Here we asked whether oncogenic signaling by KRAS and other oncogenes in lung cancer can perturb alternative splicing, possibly contributing to the mis-splicing that is so pervasive in human cancer (Climente-González et al. 2017). By expressing KRAS and other oncogenes in lung epithelial and lung adenocarcinoma cells and performing whole transcriptome analysis, we found that KRAS activation leads to differential changes in alternative RNA splicing. One alternative splicing event of note was an exon inclusion event in *MAZ*, which can regulate the expression of *KRAS* itself and *HRAS* (Ray et al. 2002). As a possible explanation for how these alternative splicing changes might occur, we identified decreased phosphorylation of SR proteins SRSF1, SRSF2, and SRSF7 in KRAS-mutant cells by LC-MS/MS. In contrast, we did not observe a change in protein abundance nor a change in gene expression of known splicing factors by KRAS variants. Thus, KRAS regulation of splicing factors occurs largely through a phospho-signaling network, underlying the importance of studying post-translational modifications.

SR proteins are critical in splice site selection in both constitutive and alternative RNA splicing as they are responsible for recognizing exon splicing enhancers (ESEs) and recruiting the spliceosome (Zhou and Fu 2013). Therefore, to modulate isoform abundance, both the abundance and activity of SR proteins in the cell, as well as the presence of ESEs in the alternate splice sites are important. SR protein activity is primarily regulated by the protein kinases Clk1 and SRPK1 (Aubol et al. 2016). In the major isoform of MAZ, exon V is preferentially selected over exon V’ likely due to the greater number of ESEs recognized by SR proteins. However, when select SR proteins are downregulated, exon V’ may be spliced in more often. In our study of KRAS-associated splicing of *MAZ*, we did not observe a change in SR protein abundances, suggesting that if the shift in splicing of *MAZ* is due to SR protein function, it is caused primarily by changes in SR phosphorylation.

By studying RNA, protein, and phosphorylation levels, we interrogated RNA splicing regulation driven by multiple oncogenes and their tumor-associated variants, identifying splicing factors and alternate exons of interest. Further studies of these alternative splicing mechanisms would open the possibility of designing treatments for mis-splicing in lung adenocarcinomas. For example, oligonucleotides can be designed and engineered to target particular splice sites in tumor cells (Havens and Hastings 2016; Levin 2019). Similarly, existing small molecule inhibitors may be used to correct splicing factor dysregulation, for example, inhibition of SRPK1 to modulate SR protein activity (Oltean et al. 2012).

Alternative isoforms in cancers also introduce neoepitopes specific to the tumor (Kahles et al. 2018). Thus, understanding oncogene-driven alternative splicing also holds implications for immune-based treatment of tumors with alterations in these oncogenes, including the design of CAR-T cells to target novel tumor neo-antigens resulting from splicing events. Further characterization of the alternate exon events associated with KRAS mutations may explain why some KRAS-mutant tumors respond to immunotherapies more favorably than others (Jeanson et al. 2019). However, our study is limited to effects *in vitro*, isolated from the immune environment, and further investigation of oncogene-driven splicing *in vivo* and in patient tumors is needed.

Additionally, although we chose to focus on exon skipping events, other forms of alternative splicing will also be critical to consider as we determine the impact to tumorigenesis and treatment (Smart et al. 2018).

## Conclusions

This work shows that diverse oncogenic signaling programs in lung cancer induce both characteristic transcriptional changes as well as alterations in alternative splicing regulation. Together with direct cis-acting somatic splicing genetic variation and trans-acting mutations in splicing factors themselves, these signaling-related splicing patterns can provide a partial explanation for the altered splicing observed in cancer. Continued large-scale studies are needed to map splicing changes to particular signaling programs and mechanistic studies are needed to better understand exactly how these programs connect to splicing. These efforts will in turn create new opportunities for therapeutic intervention in cancer via splicing modulation or immunotherapies, and both modalities may be beneficial in combination with established therapies to finally provide the durable cancer cures that have largely proven elusive.

## Figure Legends

**Supplementary Figure 1.**
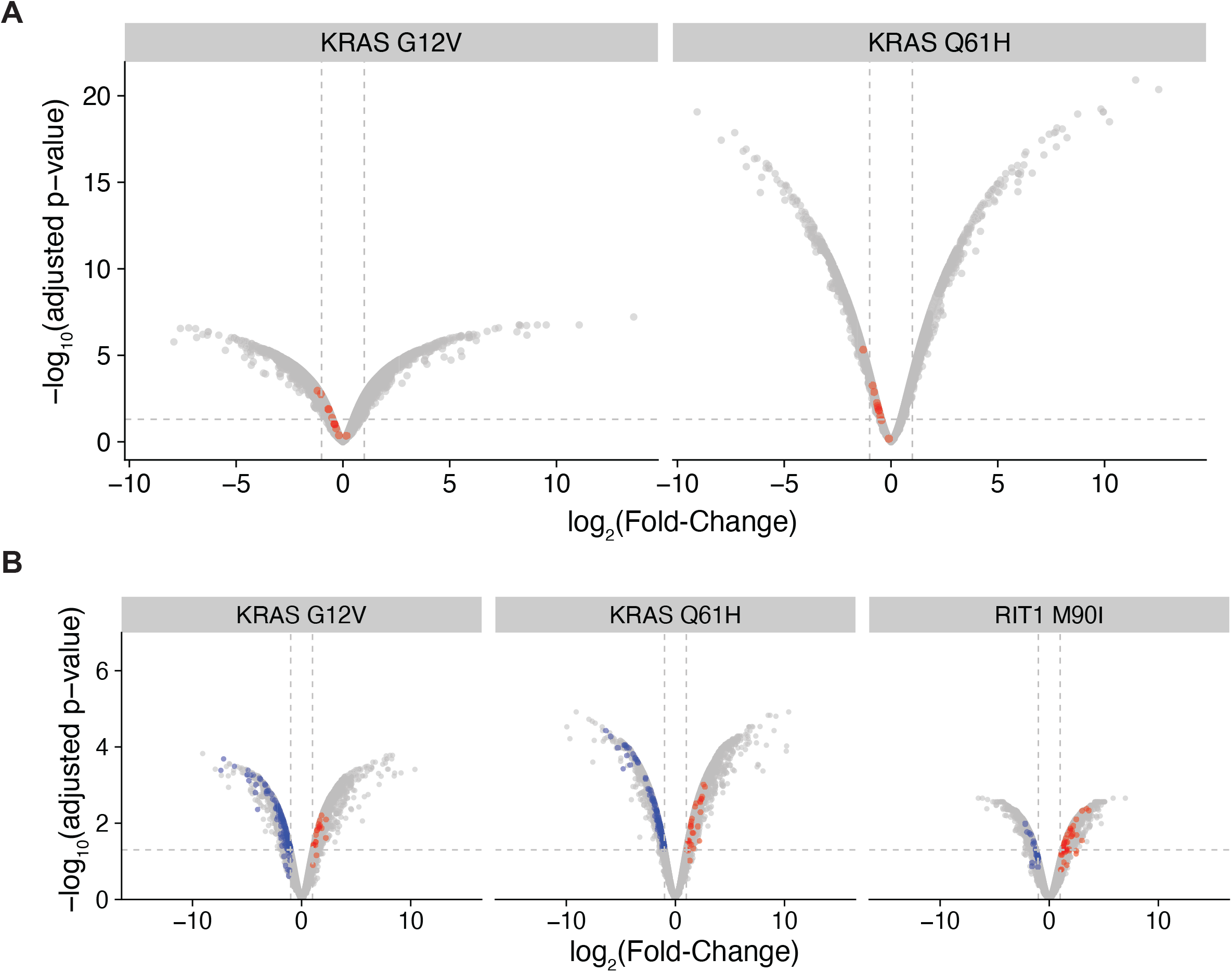
A) Volcano plot of differential protein abundance in KRAS^G12V^ and KRAS^Q61H^ cells compared to KRAS^WT^ cells. Labeled in red are SR proteins. B) Volcano plot of phosphorylation of phosphosites in KRAS^G12V^ and KRAS^Q61H^ cells compared to KRAS^WT^ cells, and RIT1^M90I^ compared to RIT1^WT^. Labeled are phosphosites on proteins in the GO RNA SPLICING gene set which are downregulated (blue) or upregulated (red).

**Supplementary Figure 2.**
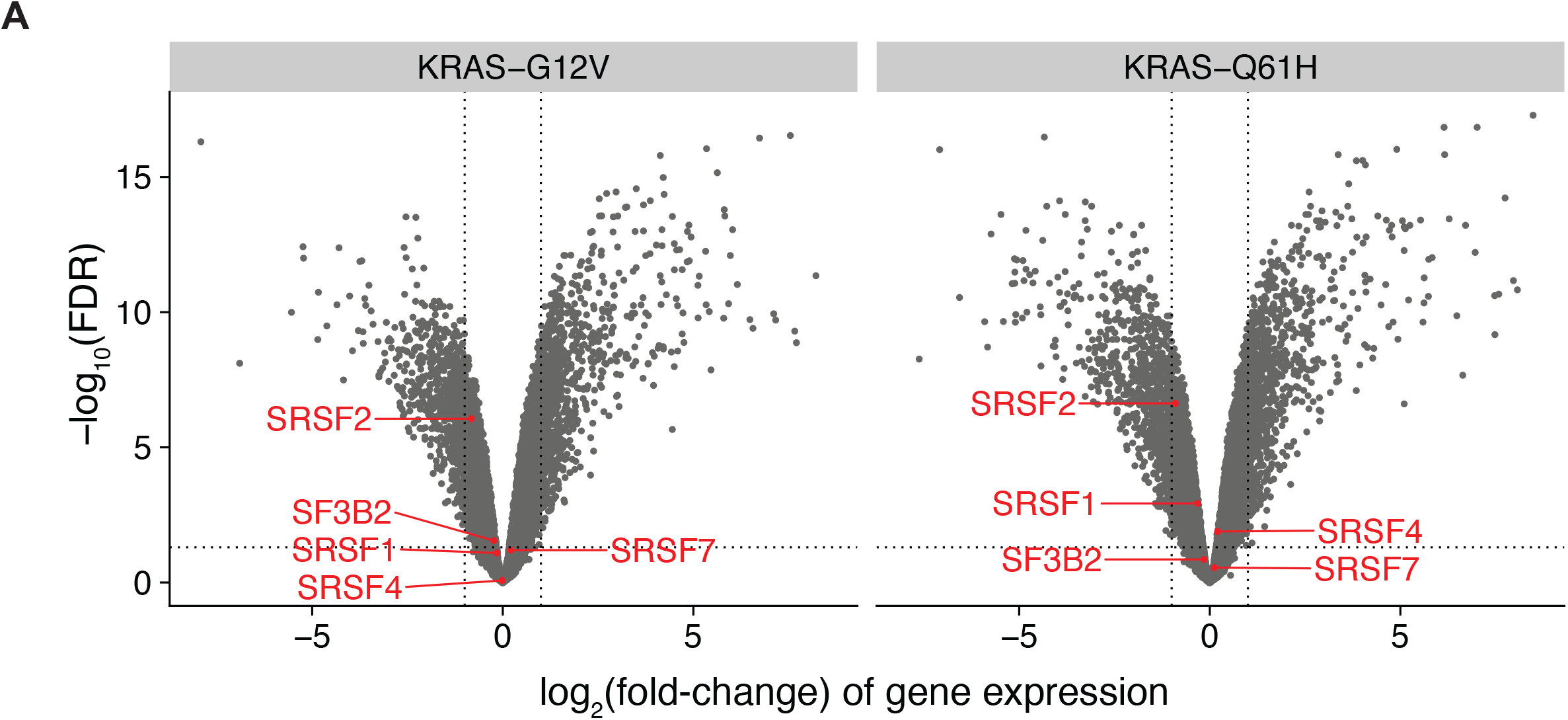
A) Volcano plot of mRNA differential expression. Labeled in red are splicing factors of interest *SF3B2, SRSF1, SRSF2, SRSF4*, and *SRSF7*. B) Change in overall gene expression compared to change in cassette exon inclusion.

**Supplementary Figure 3.**
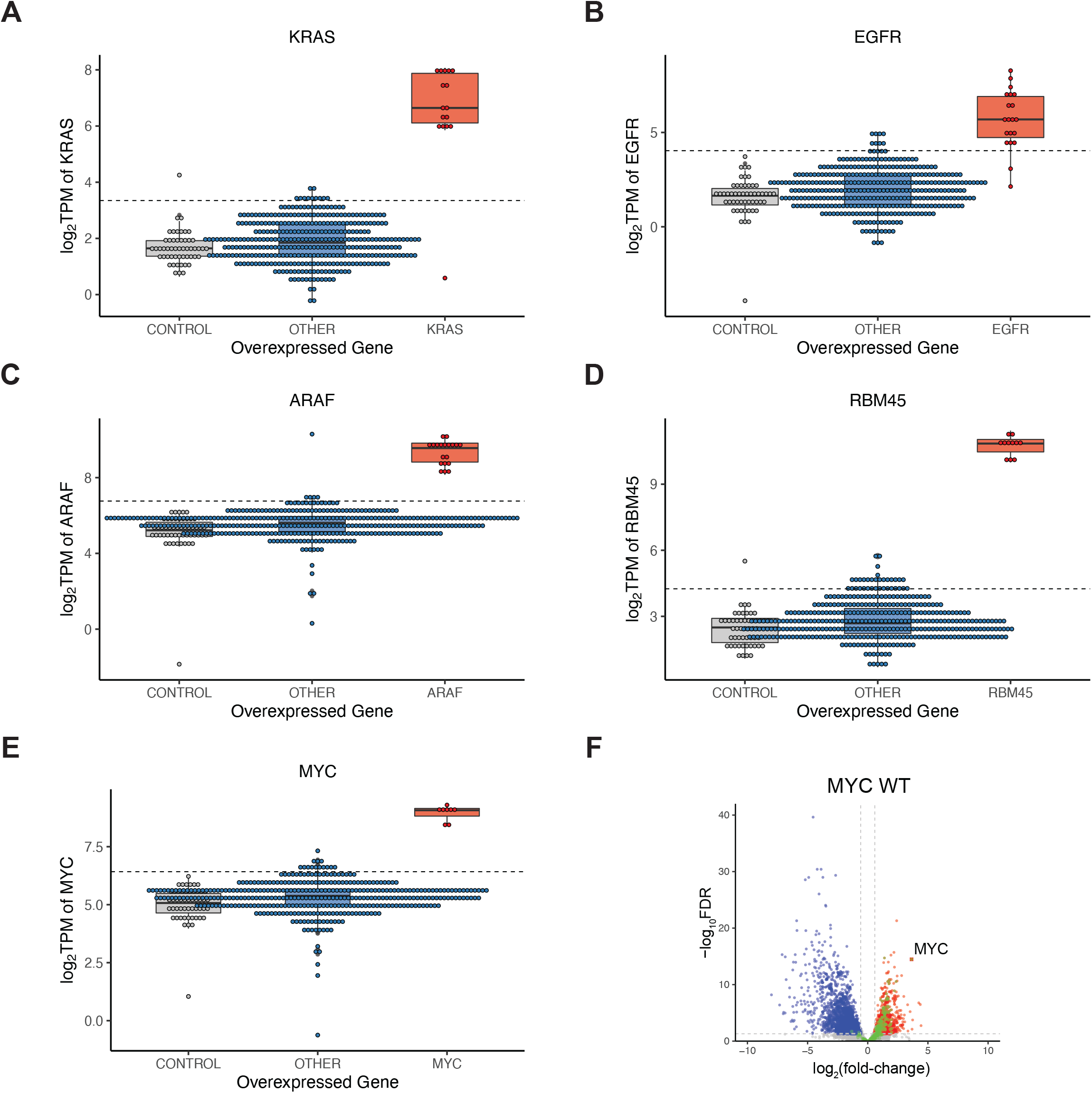
A) Volcano plot of differentially expressed genes in wild-type *MYC* overexpressing cells. Blue = downregulated transcripts. Red = upregulated transcripts. Green = transcripts in mSigDB hallmark gene sets MYC targets V1 and V2. B) mRNA expression levels of *MYC* in vector control cells (grey), cells overexpressing non-*MYC* alleles (blue), and cells overexpressing *MYC* alleles (red). C) Same as B) for *KRAS*. D) Same as B) for *EGFR*. E) Same as B) for *ARAF*. F) Same as B) for*RBM45*.

**Supplementary Figure 4.**
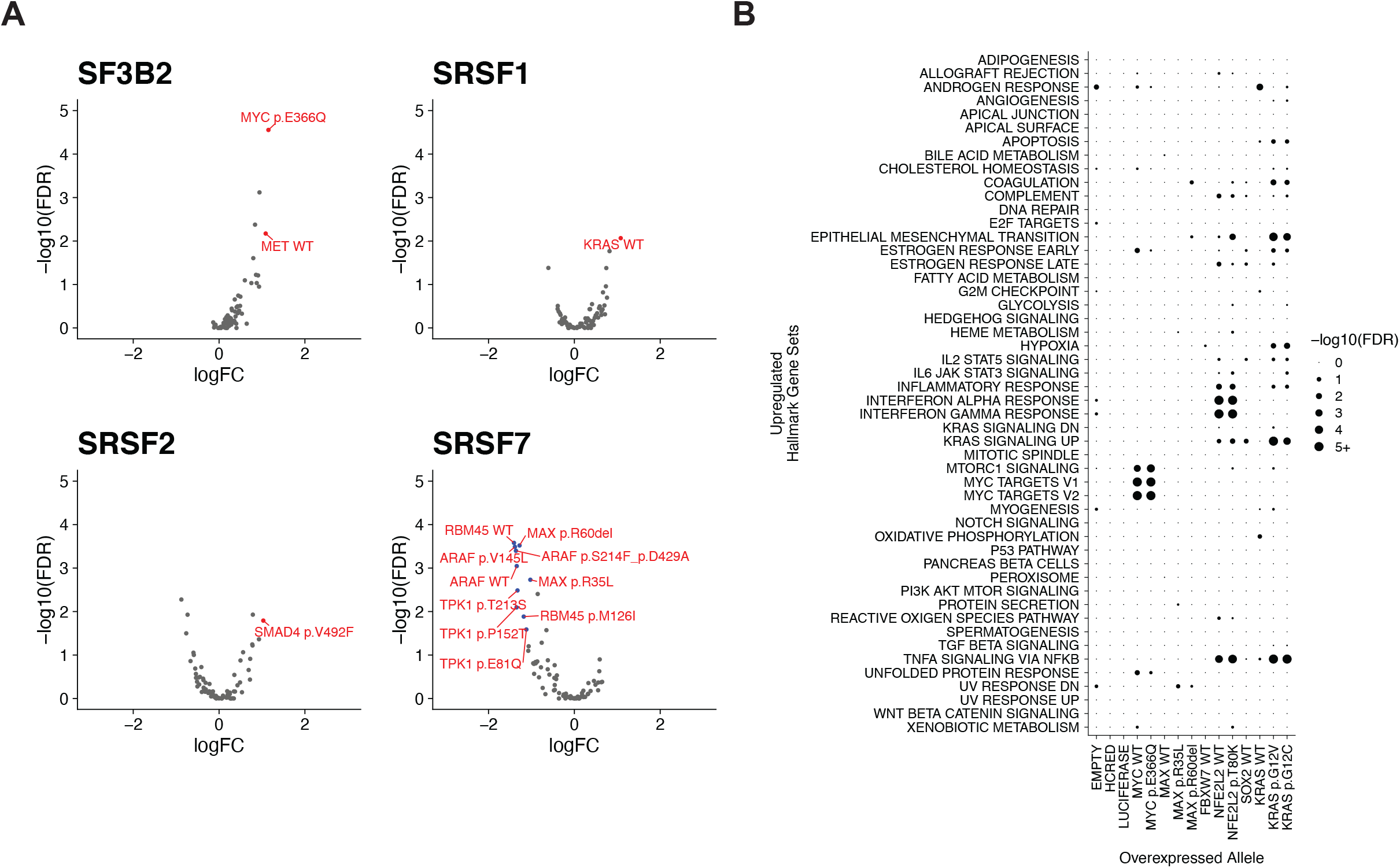
A) mRNA expression of splicing factors *SF3B1, SRSF1*, and *SRSF7*. B) Gene set enrichment analysis of downregulated transcripts in MYC, KRAS, FBXW7, and negative control cells. Hallmark gene sets from mSigDB.

**Supplementary Figure 5.**
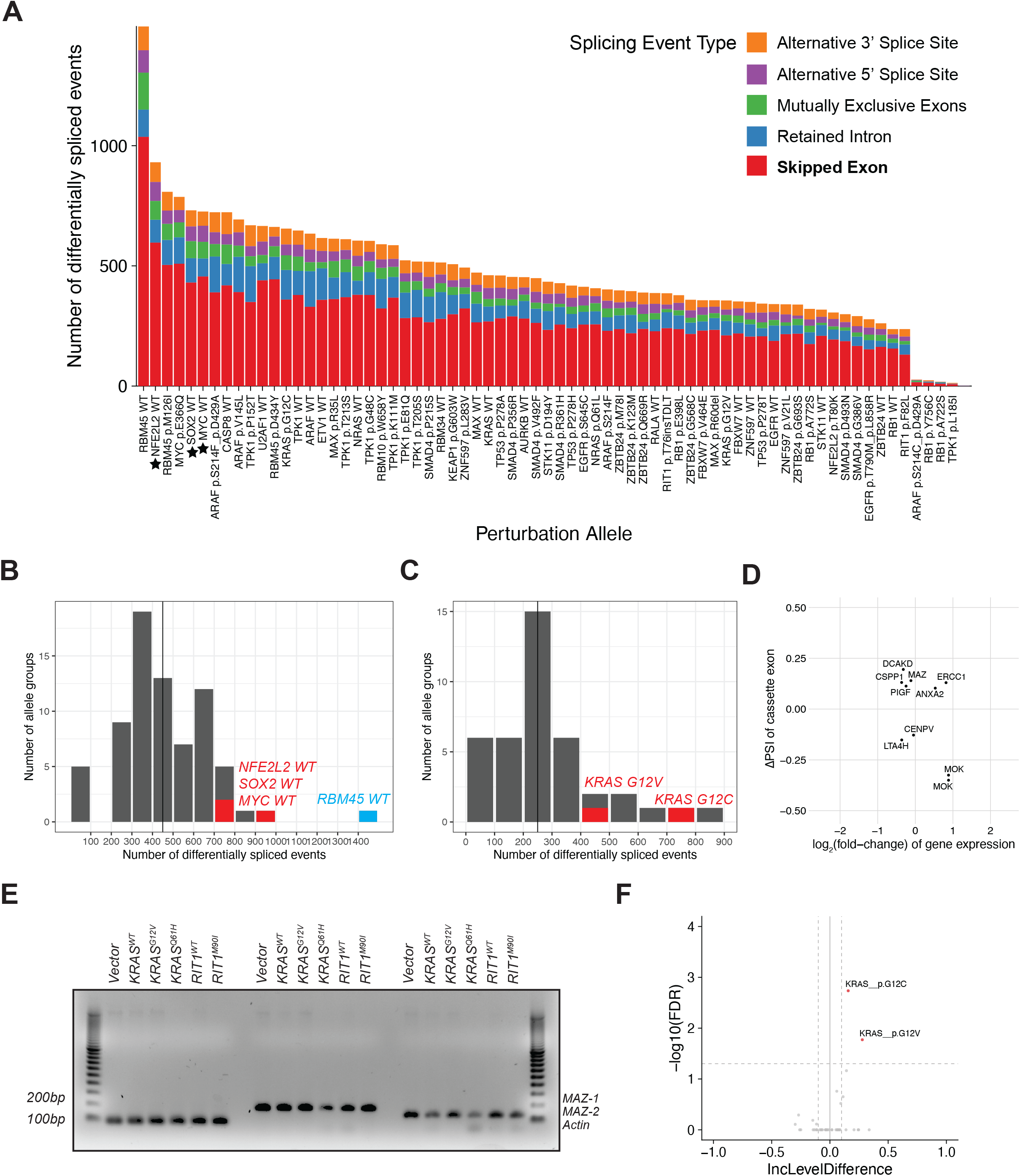
A) Number of events differentially spliced between cells with genetic perturbations and cells with vector controls. Colors represent 5 major alternative splicing event categories. Wild-type alleles of transcription factors are starred. B) Number of events differentially spliced between cells with genetic perturbations and cells with vector controls. Vertical line = mean number of events across screen. C) Number of events differentially spliced between cells with variant alleles and cells with respective wild-type alleles. Vertical line = mean number of events across screen. D) Comparing change in Percent Spliced In (ΔPSI) of cassette exons and the differential expression of the corresponding gene transcript. E) PCR gel detecting MAZ-1 and MAZ-2 isoforms in AALE cells overexpressing vector, KRAS, or RIT1 alleles. F) Differential splicing of MAZ exon V’ in variant alleles compared to respective wild-type alleles.

## Methods

### LC-MS/MS proteomics of AALE cells

Proteomic and phosphoproteomic data from AALE cells was generated by LC-MS/MS and quantified as previously described (Lo et al. 2021). Briefly, isogenic AALE cells were generated by transduction with lentiviral vectors encoding wild-type KRAS, KRAS^G12V^, KRAS^Q61H^, or wild-type RIT1 or RIT1^M90I^. Replicate lysates from each cell line were digested and labeled with TMT mass tags for quantitative mass spectrometry. Two TMT 10-plexes were used with AALE vector control lysates shared across plexes for data demultiplexing. For phosphoproteome analysis, phosphorylated peptides were enriched by IMAC column purification prior to LC-MS/MS analysis. MS data were interpreted using the Spectrum Mill software package v6.0 prerelease. All proteomic data are shared publicly in the public proteomics repository MassIVE (https://massive.ucsd.edu) and are accessible at ftp://MSV000085225@massive.ucsd.edu with username: MSV000085225 and password: oncogenic.

### RNA sequencing

RNA-seq libraries of KRAS-and RIT1-mutant AALE cells were previously described (Lo et al. 2021). Briefly, three replicates per cell line were used for RNA isolation by Direct-zol RNA mini-prep (Zymo) and libraries generated using the TruSeq RNA Library kit (Illumina). Libraries were sequenced on an Illumina NovaSeq at the Fred Hutch Genomics Shared Resource to an average coverage of 70 million 50 bp paired-end reads per sample. RNA-sequencing reads were mapped to the human genome reference hg19/GRCh37 using STAR 2.5.3a. Data is publicly available at the NCBI Gene Expression Omnibus database with accession number GSE146479.

For the A549 large-scale splicing screen, A549 lung cancer cells were transduced at high multiplicity-of-infection in 384 well plates using pLX317-ORF constructs as previously described with 4-6 biological replicates per ORF (Berger et al. 2016). Cells were not selected with puromycin but parallel plates were treated with puromycin or left untreated and then cell viability was determined using CellTiterGlo reagent (Promega) for calculation of infection efficiency. 96 hours post-transduction, cells were lysed using TCL buffer (Qiagen) and lysates were stored at -80 degrees. We adapted library preparation protocols previously developed for single cell RNA sequencing (Trombetta et al. 2014). RNA was isolated from 9 µl lysate and 1 µl of ERCC 1:1000 Spike in control (Thermo Fisher Scientific) per sample using RNA SPRI beads (Agencourt) in lo-bind microcentrifuge tubes (Eppendorf). Beads were washed several times in 80% ethanol before drying and used for cDNA synthesis. First-strand cDNA synthesis was performed with the Smart-seq v4 Ultra Low Input kit (Takara) using the 3’ Smart-seq CDS Primer IIA and Smart-seq V4 oligonucleotide. Next, whole transcriptome amplification and clean-up was performed using the Kapa HiFi HotStart kit with the following thermal cycling protocol: 3min 98°C; 20 cycles of 15s at 98°C, 15s 67°C, 6min at 72°C; final extension of 5min at 72°C. DNA was then cleaned up using DNA SPRI beads (Agencourt). Individual samples were inspected by Qubit quantitation and TapeStation (Agilent) analysis and diluted to 0.1 to 0.2 ng/µl prior to library construction using the Nextera XT sequencing kit (Illumina). Final libraries were pooled and sequenced on an Illumina HiSeq (Northwest Genomics Center) to an average coverage of 10M 75 bp paired-end reads per replicate. RNA-sequencing reads were mapped to the human genome reference hg19/GRCh37 using STAR 2.5.3a.

### Differential gene expression analysis

Transcripts were quantified with featureCounts from the Subread package v.1.5.3 (Liao et al. 2014) using RefSeq gene annotations. Differential expression was then determined using edgeR v.3.30.3 (Robinson et al. 2009), normalizing transcript counts by library size and RNA composition scale factors computed by using trimmed mean of M-values (TMM) between sample pairs (Robinson and Oshlack 2010). Whole transcriptome profiles were quality checked for sufficient RNA-seq coverage. Samples were filtered for successful expression of the lentiviral vector, determined by a 2 standard deviation increase in expression of the target gene in the sample compared to negative controls. Gene transcripts were also filtered for sufficient detection across the experiment as determined by a mean logCPM > 0.1 for each gene. After applying quality control filters, the A549 dataset included 374 transcriptome profiles each quantifying 12543 genes. Analysis pipeline is available as a Snakemake workflow at https://github.com/bergerbio/RNA-splicing-screen.

### Differential splicing analysis

After alignment of RNA-seq reads by STAR, alternative splicing events were identified and quantified by rMATS-turbo 4.1.1 using gene transcript annotations from gencode v.19 (Shen et al. 2014). Both reads spanning exon junctions and reads covering single exons were used for splicing quantification. Analysis pipeline is available as a Snakemake workflow along with the in-house R scripts used to aggregate rMATS results and perform additional analyses and interpretations (https://github.com/bergerbio/RNA-splicing-screen).

### Gene Set Analysis

Analysis of enrichment of KRAS signaling in differential RNA expression profiles was performed in R with GOseq (Young et al. 2010). KRAS signaling gene sets were taken from MSigDB hallmark gene sets (Liberzon et al. 2015).

### Quantitative reverse transcription polymerase chain reaction (qRT-PCR)

Total RNA was extracted using Direct-zol RNA Miniprep plus (Zymo Research) from each biological replicate of AALE cell lines overexpressing KRAS or RIT1 alleles or control vector. From the extracted RNA, 1 µg was reverse transcribed into cDNA using SuperScript IV First-Strand Synthesis (Invitrogen). Quantitative RT-PCR reactions were set up in technical triplicates with BioRad iQ SYBR Green kit and analyzed on a BioRad CFX384 Real-Time System. PCR primers were designed to detect MAZ isoforms without alternate exon V’ (Forward: ATG GGA GGC AGC TTT CGC; Reverse: TCA CCA GTA CCT TTG TTG CA) and with exon V’ (Forward: AGCTCTGCAACAAAGGCTTC; Reverse: GGGCAGGGGTCTTGCA). PCR products were confirmed to be specific by molecular weight analysis via gel electrophoresis. Quantification of isoforms in experimental samples were normalized against vector control samples and relative quantification of MAZ isoforms 1 and 2 were calculated with the Livak method.

## Declarations

### Ethics approval and consent to participate

Not applicable

### Consent for publication

Not applicable

### Availability of data and materials

Proteomic data analyzed in the current study are available in the public proteomics repository MassIVE (https://massive.ucsd.edu) and are accessible at ftp://MSV000085225@massive.ucsd.edu with username: MSV000085225 and password: oncogenic. RNA-seq data generated and analyzed in the current study are available in the Gene Expression Omnibus (GEO) repository and are accessible at GSE146479, GSE183670, and [GEO Accession number will be provided before review/publication].

### Competing interests

AHB reports consulting fees from Puma Biotechnology.

### Funding

This research was funded, in part, through the National Cancer Institute (NCI) R37 CA25205 to A.H.B., NIH/NCI Cancer Center Support grant P30 CA015704, and NSF IGERT DGE-1258485 to A.L.

## Authors’ contributions

A.H.B. designed and supervised the study and provided study resources. A.L., M.M., and A.H.B. performed experiments. A.L. performed all computational analyses of the proteomics, transcriptomics, and splicing. A.L. and A.H.B. wrote the manuscript. All authors read and approved of the final manuscript.

## Acknowledgements

We thank Rob Bradley, Daniela Witten, Cole Trapnell, David MacPherson, and Berger and MacPherson lab members for critical advice and discussion. We thank Angela Brooks, Xiaoyun Wu, Jesse Boehm, and the Broad Institute Genetic Perturbation Platform for our previous collaboration in generating the large-scale screening platform. We thank Fil Mundt, Philip Merins, and Steve Carr and the Broad Institute Proteomics Platform for our prior collaboration in generating the proteomic data. We thank the Northwest Genomics Center and Fred Hutch Genomics Shared Resource for generation of RNA sequencing data.

